# Enhanced reward coding and condition-independent dynamics in optogenetically identified corticostriatal neurons in monkeys

**DOI:** 10.1101/2025.04.21.649760

**Authors:** Adi Hovav-Lixenberg, Yirat Henshke, Tirzah Kreisel, Eran Lottem, Mati Joshua

## Abstract

The basal ganglia are considered to be the site where cortical sensorimotor and dopaminergic reward information interact to potentiate and select actions. This had led to the assumption that cortical inputs to the basal ganglia encode sensorimotor states rather than reward or choice signals. We tested this hypothesis by studying the coding properties of neurons in the frontal eye field of monkeys that were optogenetically identified as being connected to the basal ganglia. We found that neurons already contained information about expected rewards and selected actions. Further, the reward and condition-independent modulations were stronger in connected neurons than in other neurons in the same area. These findings indicate that reward, choice, and sensorimotor information are integrated already in the inputs to the basal ganglia, implying that the basal ganglia play a role in manipulating rather than generating reward and choice signals.

## Introduction

One of the widespread assumptions about basal ganglia function is that the striatum integrates reward information from dopaminergic inputs with cortical information to implement reinforcement learning algorithms for action selection^1–3^. Dopaminergic input is hypothesized to provide the reward prediction error signal^4–6^ that acts on state information from the cortex. Extensive testing has for the most part supported the coding of prediction error by dopamine neurons, though some variations have emerged^7–9^. Unlike the dopaminergic input that has been studied comprehensively, the content of the cortical input to the basal ganglia remains less understood^10,11^. Cortical neurons have been shown to integrate reward, choice and sensorimotor information^12–17^, but neurons projecting to the basal ganglia may not contain reward signals. Further, other signals transmitted from the cortex to the basal ganglia, aside from reward-related signals, remain poorly characterized.

To study the content of cortico-striatal signaling, we focused on the monkey eye movement system. Due to the experimental control it affords and similarity to humans in neural circuitry and movement repertoire, this system is among the most frequently studied and best understood^18,19^. The frontal eye field in the frontal cortex is connected to the caudate nucleus in the striatum^20,21^, and both areas are causally linked to eye movements^22,23^. We aimed to characterize the information transmitted from the FEF to the caudate.

In earlier research protocols, input identification in monkeys used antidromic electrical stimulation, where axons of neurons were stimulated at the target site and detected at the source^10,24^. However, this method has a number of shortcomings; most prominently, the low probability of successfully simultaneously stimulating and recording projection neurons. Viruses that code channelrhodopsin and are transported retrogradely from the injection site (e.g., the caudate) to the source (e.g., FEF) provide an effective alternative^25,26^. The input neurons can be identified by their response to light stimulation^27–29^, a method termed opto-tagging. We opto-tagged neurons projecting from the FEF to the caudate and recorded their activity during eye movement tasks where reward size was manipulated. We found that neurons connected to the caudate already encoded strong reward and choice information and exhibited more pronounced condition-independent modulations than other neurons, thus suggesting that the integration of reward, choice and sensorimotor occurs already in the inputs to the basal ganglia. These results, along with our previous findings of a sharp increase in temporal complexity^30^ and the signal-to-noise ratio of neurons in basal ganglia output^31^ provide evidence that the basal ganglia process reward and choice information rather than generate these signals.

## Results

### Reward impacts monkeys’ behavior in eye movement tasks

The monkeys were engaged in three eye movement tasks in which we manipulated the reward size (Fig. 1A). In these tasks, the monkeys were presented with color-coded targets that indicated whether the monkeys would receive a large or small reward at the end of the trial. The monkeys observed a colored target that appeared in a central (the pursuit task) or a peripheral location (the saccade task). After a variable delay, the monkeys were instructed to move to the target (Go). On the saccade trials, the monkeys moved their eyes ballistically to the eccentric location (Fig. 1A and B, left), whereas on the pursuit trials the monkeys smoothly followed a continuously moving target (Fig. 1A and B, middle). Saccade and pursuit movements were directed to one of the four cardinal directions. In the choice task, two targets appeared and the monkeys had to select one (Fig. 1A and B right). At the end of each trial, the monkeys received either a small or large reward based on the color of the single or selected target. We refer to the epoch during which the monkeys observed the cues but were not allowed to move as the *cue epoch*, the epoch during which they moved their eyes to the colored target as the *movement epoch*, and the epoch during which they received the reward as the *outcome epoch*.

**Figure 1:**
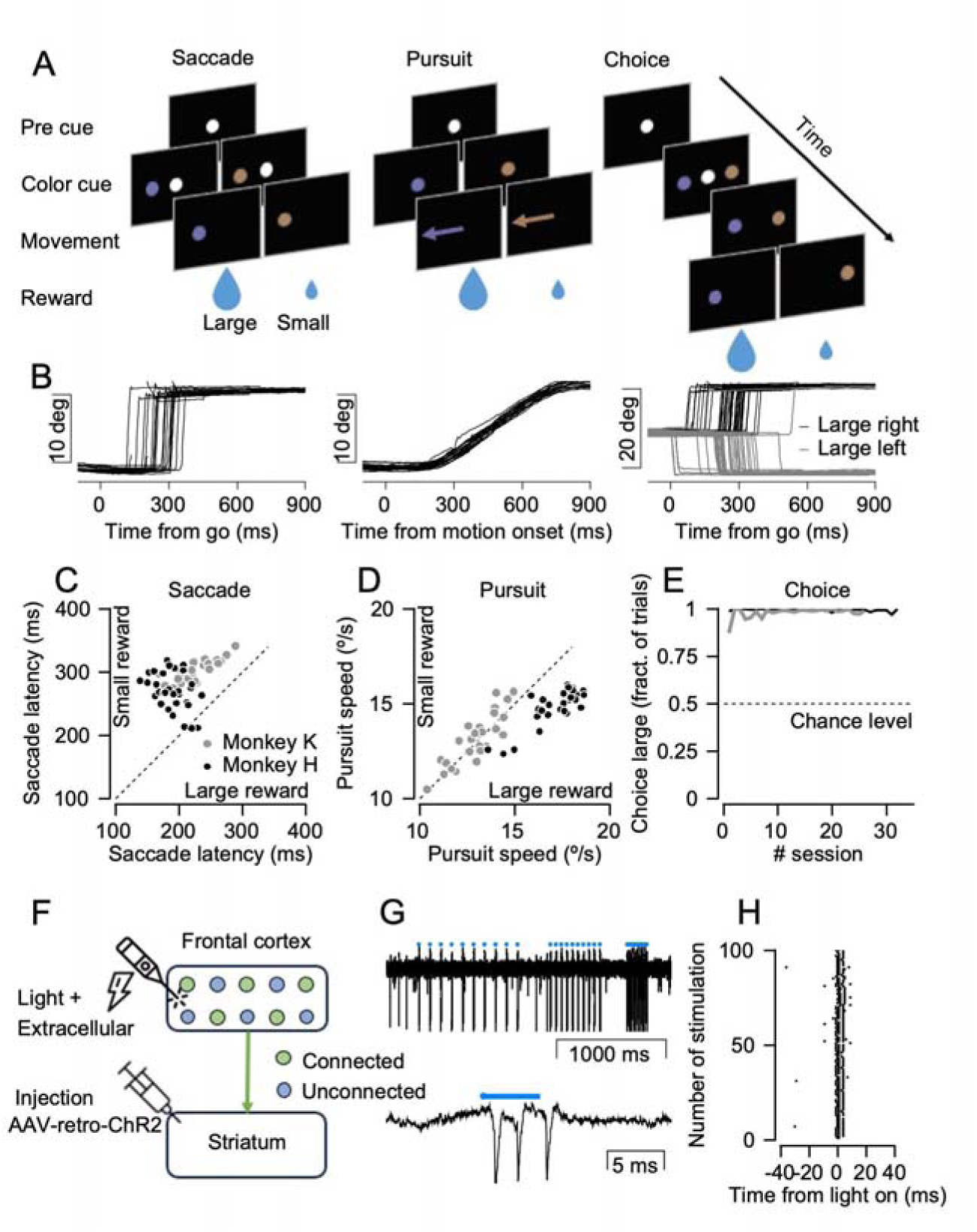
Task behavior and opto-tagging of FEF neurons. **A.** Schematics of the three tasks. The series of black squares represent the consecutive visual stimuli that were presented on the screen. Dots show stationary targets and arrows indicate moving targets. The size of the drop corresponds to the reward size. **B.** Example of traces of the eye position in the horizontal direction; positive and negative values correspond to movement to the right and left. Each thin line shows a single trial; traces correspond to the task schematics above them. **C-E:** Day-by-day analysis of the behavior of the monkeys on the saccade (**C**) and pursuit tasks (**D**). Each dot shows the saccade latency (**C**) or average eye velocity 200 -250 ms after motion onset (**D**) averaged across task condition and trials from a single day. Horizontal and vertical axis show the large and small reward conditions. **E.** Fraction of trials in which the large reward was selected as a function of the recording session. Only days in which we recorded neural data are included. **F.** Schematics showing the injection in the striatum and recording in the FEF. **G**. Example of an extracellular recording during light stimulation. Blue dots and the blue line show the time of the stimulation (5 ms duration). **H.** Raster of the response of a single neuron to 100 light stimulations. Each dot shows the time of a single spike aligned to the time of the stimulation. Only trials from the 10 Hz stimulation are shown to be able to depict the pre-stimulus activity without confounding with the responses to the previous stimulation.

The monkeys’ behavior indicated that they associated the colored target with the upcoming reward. On the saccade tasks, the monkeys’ gaze moved earlier in the large reward target (Fig. 1C, p=8*10^-5 and 10^-6 for monkeys H and K, Wilcoxon signed-rank test). On the pursuit trials, the eye velocity at the movement initiation of one monkey was consistently faster for the large reward (Fig. 1D, eye velocity at 200-250 ms after target motion onset, p = 8*10^-7 and p =0.98 monkeys H and K, Wilcoxon signed-rank test). The monkeys’ preferences were confirmed on a choice task in which both monkeys almost always selected the target that was associated with the larger reward (Fig. 1E p = 5*10^-7 and p=8*10^-6 for monkeys H and K, Wilcoxon signed-rank test for difference from a chance level of 0.5), demonstrating that the monkeys associated color and reward size.

### Opto-tagging of neurons in the FEF connected to the striatum

To identify neurons in the FEF connected to the basal ganglia, we injected an AAVretro-ChR2 virus^25^ into the eye movement areas of the caudate (Fig. 1F and Fig. S1). The virus is transported retrogradely from the axons in the caudate to the cell bodies in the cortex. As a result, in the cortex, only cortico-striatal projection neurons express ChR2. We confirmed the injection location over the course of several days during which we injected a solution with Mn+2 and imaged the injection site with an MRI scan (Fig. S1). We waited four weeks for viral expression and then on each recording day we lowered an optrode with 4 closely spaced recording contacts attached to a fiber for light stimulation (Thomas Recordings) into the FEF (Fig. S2). We searched for neurons that responded consistently with short latencies (see Methods) to brief light stimulations (durations of 5 ms at 10, 20 and 50 Hz). Figure 1G shows an example of the extracellular activity in one contact of the optrodes while we applied the light stimulation. The recorded neuron fired sparsely before the stimulation and very consistently with a very short latency during the stimulation (Fig. 1G, H). Based on the response to stimulation (see Methods) we categorized the neurons into (1) light responding neurons that were highly likely to be cortico-striatal projection neurons, (2) neurons that were recorded in the same location as the responding neurons but did not respond to the stimulation, or their response was inconsistent, and (3) neurons in which none of the neurons in the recording session responded consistently to the stimulation. A response to light was considered strong evidence for the cell to be a cortico-striatal neuron, but a lack of response was harder to interpret. For this reason, we classified the neurons into three groups: *tagged, untagged* and *area-untagged*. We performed an offline analysis to confirm the consistency of the extracellular waveforms of the connected neurons across the stimulation and behavioral sessions (Fig. S3 and Methods).

### Factoring the response of single neurons to the experimental variables

To study the coding properties of the neurons, we factored the single-neuron activity into the direction, reward and condition-independent components. Figure 2A-E shows the response of an example neuron to the appearance of an eccentric color cue in the saccade task. Averaging across either the reward or direction conditions (Fig. 2C and D) showed that the neuron responded to the direction of movement and to the reward conditions. Averaging across all conditions showed that the neuron exhibited overall temporal modulations which were not dependent on condition (Fig. 2E). To quantify the contribution of these different experimental variables to the overall activity, we calculated the partial *ω*^2^ effect size (*ω*^2^)^31,32^.

**Figure 2:**
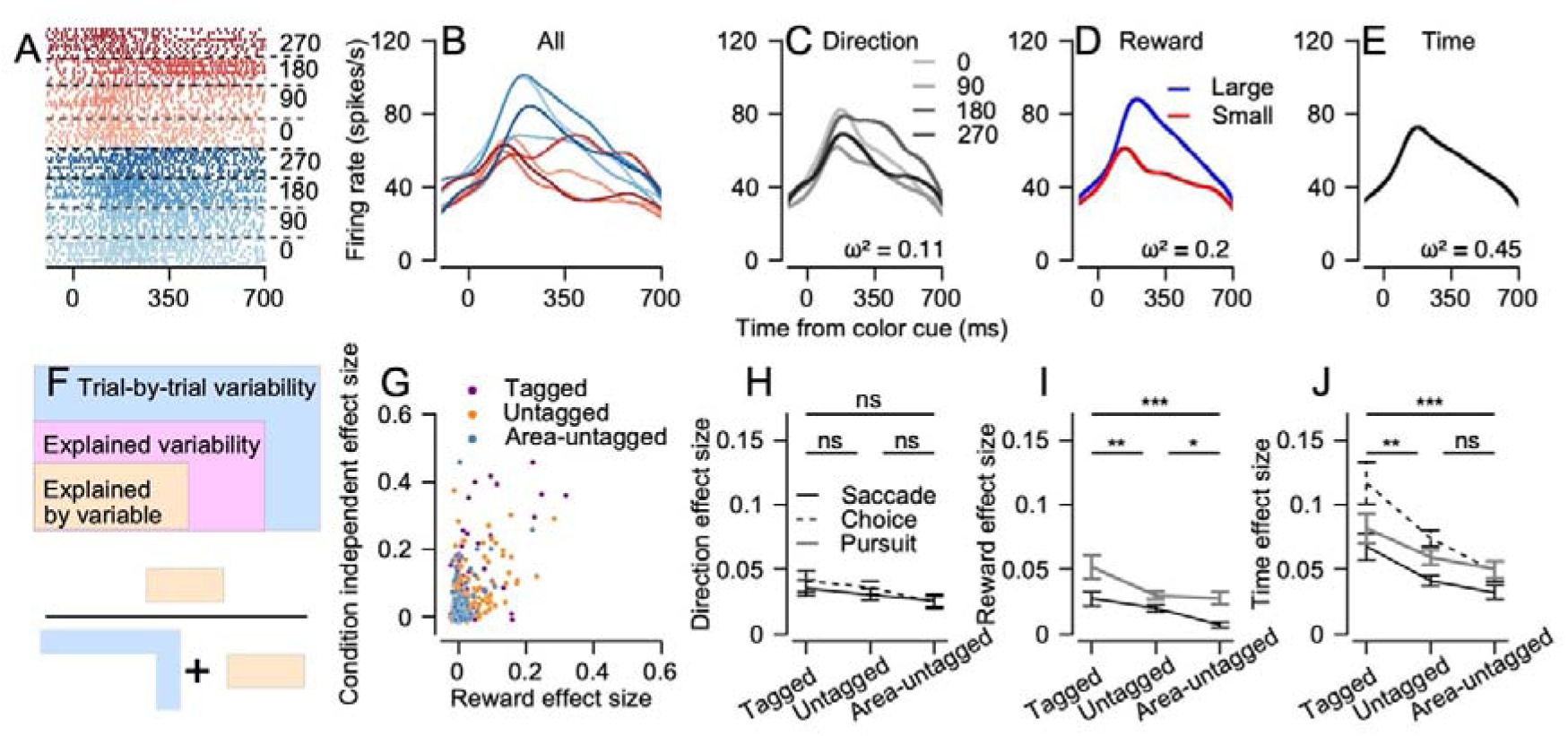
Effect size analysis during the cue epoch. **A-E.** Example of responses of one neuron during the cue epoch of the saccade task. Blue and red traces correspond to the large and small reward conditions. The luminance (light to dark) corresponds to the direction of movement. **A** and **B** show the raster and PSTH for all conditions separately**. C-E** show traces in which the PSTHs were averaged across the reward (**C**), direction (**D**) and all (**E**) conditions. The ω^2^ in the figure shows the effect size associated with each of the variables. **F**. Schematics demonstrating the calculation of the effect size. The outer square represents the total trial-by-trial variability, the intermediate square shows the variability that can be explained by the experimental variables and the inner square shows the variability that is explained by one specific variable. The lower schematics demonstrate the calculation of the variability components. **G.** Example of the distribution of effect size. Each dot shows the effect size for the reward (horizontal) and time (vertical) variables for a single neuron. Different colors correspond to the different populations. **H-J**. Summary of the effect sizes during the cue for the three populations. Different plots show the effect size for the direction (H), reward (I) and time (J) variables. Vertical values show the averages and error bars show the SEM. Lines above show the results of the permutation test for average differences across all tasks (ns – not significant, * p < 0.05, ** p<0.01, *** p< 0.001).

Specifically, *ω*^2^ is calculated as the trial-by-trial variability explained by an experimental variable divided by the sum of this same variability and a variability that is not explained by the variables (noise). In other words, it quantifies the trial-by-trial signal-to-noise of a neuron (Fig. 2F). *ω*^2^ is distributed around zero (unbiased) when a neuron does not respond to the experimental variable and has a value of 1 when all the trial-by-trial variability is explained by an experimental variable. Intermediate values indicate how much of the neuron’s activity can be accounted for by a specific variable in comparison to the trial-by-trial variability that cannot be explained by any of the variables. To calculate the effect size, we used an ANOVA model for each neuron, with reward size, direction of movement, and the time bin within the trial as variables (see Methods, e.g., values in Fig. 2C–E). The direction effect size estimates the magnitude of the neuron’s direction tuning, the reward effect size estimates reward-size related modulation, and the time effect size estimates modulations that are condition-independent, i.e., overall average modulations across all conditions (e.g., Fig. 2E).

The results of the effect size analysis confirmed findings from previous studies: (1) The direction and reward effect sizes were significantly different from zero (Fig. 2H–I) as expected from neurons in the FEF^13,15^, (2) The effect size was widely distributed with most of the neurons having small values close to zero but some exhibiting large effect sizes (e.g., Fig. 2G) demonstrating the typical long-tailed coding of task parameters^33^, (3) The time effect size tended to be the largest (Fig. 2J and see for example the comparison in Fig. 2G between reward and condition-independent effect sizes), consistent with previous studies reporting that condition-independent modulations can account for the largest share of the population activity^14,34^.

### Tagged neurons had the largest time and reward effect sizes during the cue epoch

We compared the effect sizes of the three populations of neurons. On all three tasks, the average time effect size of the tagged neurons was larger than in the untagged and the area-untagged populations (Fig. 2J). In the tasks where we manipulated the upcoming reward size (saccade and pursuit), the reward effect size was larger in the tagged neurons (Fig. 2I). On the tasks where the visual eccentric target provided information about the direction of the upcoming movement (saccade and choice), the direction effect size was not significantly different across populations (Fig. 2H). Thus, the effect size analysis of the cue period indicated that reward and time modulations were the most strongly observed by inputs to the basal ganglia.

### Larger time and reward effect sizes in the tagged neurons persisted into the movement epoch

The FEF demonstrated coding of multiple experimental variables during the movement epoch as well. Figure 3A shows an example neuron that coded multiple experimental variables during the saccade tasks around the time of the saccade. This neuron responded with a burst of activity around the time of the saccade. Interestingly, its response was stronger in the small reward condition (red versus blue lines in Fig. 3B,D). Later we return to an analysis that quantified whether and when reward size was coded by an increase or decrease in activity. The same neuron also responded differently for different directions of movement (Fig. 3C) and the overall modulations were not averaged out across conditions (Fig. 3E,). This coding of multiple components was reflected in the positive values of the effect size.

**Figure 3:**
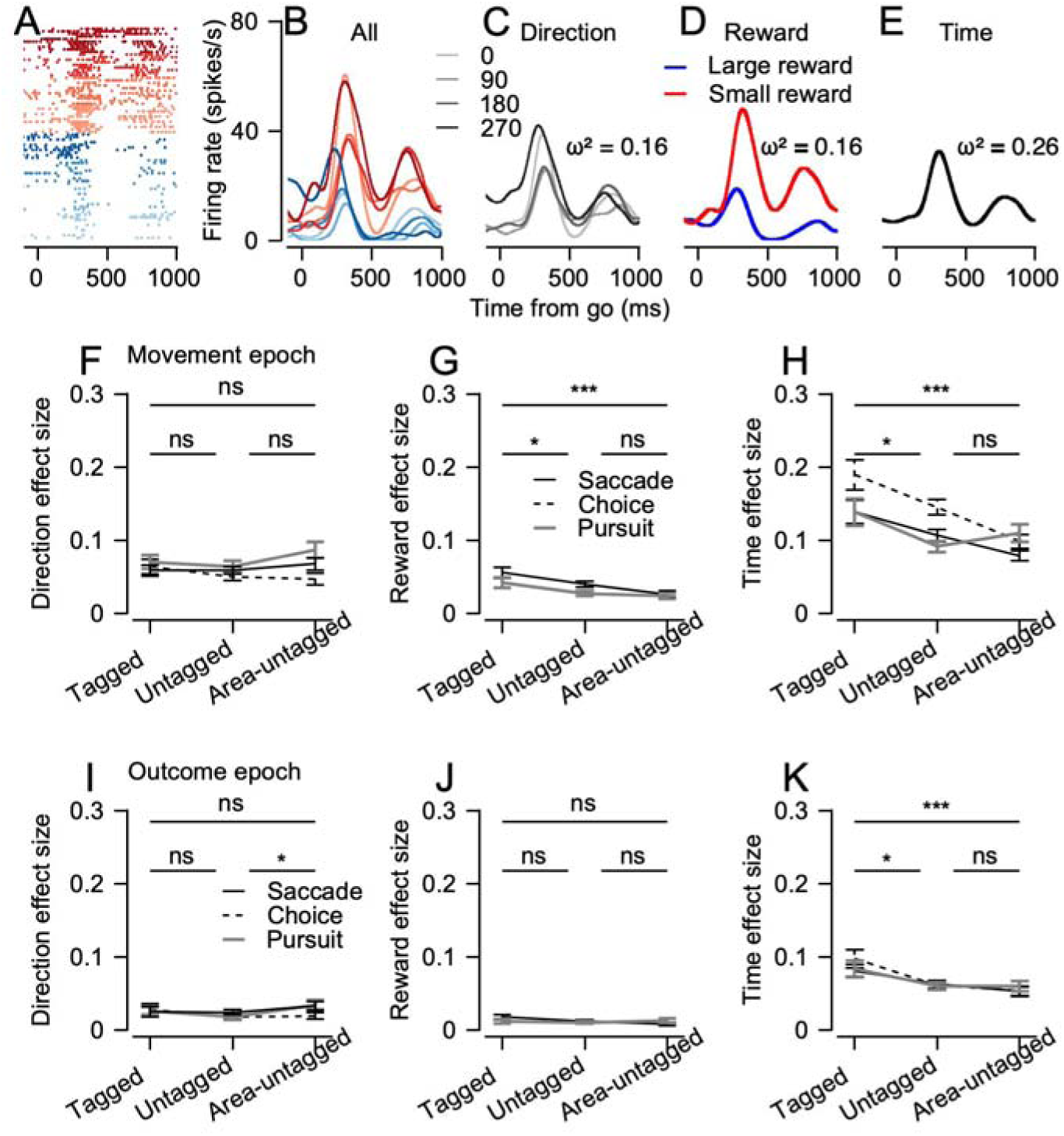
Effect size analysis during the movement epoch. **A.** Example of responses of a neuron during the movement epoch of the saccade task. Blue and red traces correspond to the large and small reward conditions. The luminance (light to dark) corresponds to the direction of movement. **A** and **B** show the raster and PSTH for all conditions separately. **C-E** show traces in which the PSTHs were averaged across the reward (**C**), direction (**D**) and all (**E**) conditions. The ω^2^ in the figure shows the effect size associated with each of the variables. **F-K**. Summary of the effect sizes during movement (**F-H**) and outcome (**I-K**) for the three populations and tasks. Different plots show the effect size for the direction (**F,I**), reward (**G,J**) and time variables (**H,K**). Vertical values show the averages and error bars show the SEM. Lines above show results of the permutation test for average differences across all tasks (ns – not significant, * p < 0.05, ** p<0.01, *** p< 0.001).

We then compared the effect sizes across populations. As was the case for the cue, the tagged neurons had the largest reward and time effect sizes (Fig. 3G and H). The direction effect size was not significantly different across populations. Thus, analysis of the movement epoch further demonstrated the coding of reward information by inputs to the basal ganglia and the larger condition-independent modulations of these inputs. We also calculated the effect sizes during the outcome (Fig. 3I-K). This analysis further supported the larger time effect size by the tagged neurons (Fig. 3K). We did not find any difference in the coding of reward at the outcome (Fig. 3J); however, the overall reward modulations were very small at this time. In fact, the cue and movement epoch reward effect sizes were larger than the reward effect size for the outcome (p =9*10^-9 and p=10^-33, outcome vs. cue and outcome vs. movement, permutation test, see Methods). Thus, the FEF neurons were modulated more strongly by the color cues that induced reward expectation than by the reward itself.

In addition to the effect size differences between the populations, we also found that the baseline firing rate defined by the pre-cue activity (Fig. 1A pre-cue) was larger for the tagged neurons. Even when taking this firing rate baseline into account, the time and reward effect sizes were larger for the connected neurons (Fig. S4).

### Beyond effect size: taking into account the sign of the modulation

The effect size analysis has the advantage that it factors activity into the experimental variables prompting comparison between populations. However, the effect size does not indicate whether the modulations are a result of an increase or decrease in rate. For example, the neurons shown in Figure 2A and 3A responded differently to the large and small reward conditions and therefore had a positive reward effect size; however, their responses were qualitatively different since one neuron responded more strongly to the large reward whereas the other responded more strongly to the small reward. Another limitation of effect size is that the nature of the interaction between variables can be difficult to interpret. For example, interaction between reward and direction could result from either reward potentiation or attenuation of direction tuning or other patterns.

To further study if and how reward is processed in the different populations, we performed the next analyses by taking the sign of the rate modulation into account. We aim to test whether (1) the sign of the reward modulation is consistent across neurons, and (2) how reward modulations interact with the direction tuning curves.

### Reward potentiated directional tuning in the cue epoch the most strongly for tagged neurons

To study how reward modulated the coding of direction, we first tested whether each neuron was directionally tuned (ANOVA for direction, p < 0.05) and then calculated the preferred direction (PD, see Methods). We then calculated the average activity in the PD and the opposite direction (Null) for the large and small reward conditions. Directional tuning was enhanced by the reward, as indicated by the larger responses in the PD for the large versus small reward condition (Fig. 4A-C). Quantification of reward modulation in the PD at the single-neuron level showed that neurons across all the populations tended to respond more strongly in the large reward condition (Fig. 4E-G). The effect of the reward in the PD was the largest in the tagged neurons (Fig. 4D, solid black line): the difference in rate between the large and small rewards was significantly greater for the tagged neurons than for the other groups (p=0.02 and p=0.002 for tagged versus untagged and tagged versus area-untagged, Wilcoxon rank-sum test). Thus, the directional tuning of the visual responses of inputs to the basal ganglia was already strongly potentiated by reward. This potentiation of visual responses by reward is akin to reports of modulations of spatial attention^35,36^, suggesting that inputs to the basal ganglia already carry spatial attention information.

**Figure 4:**
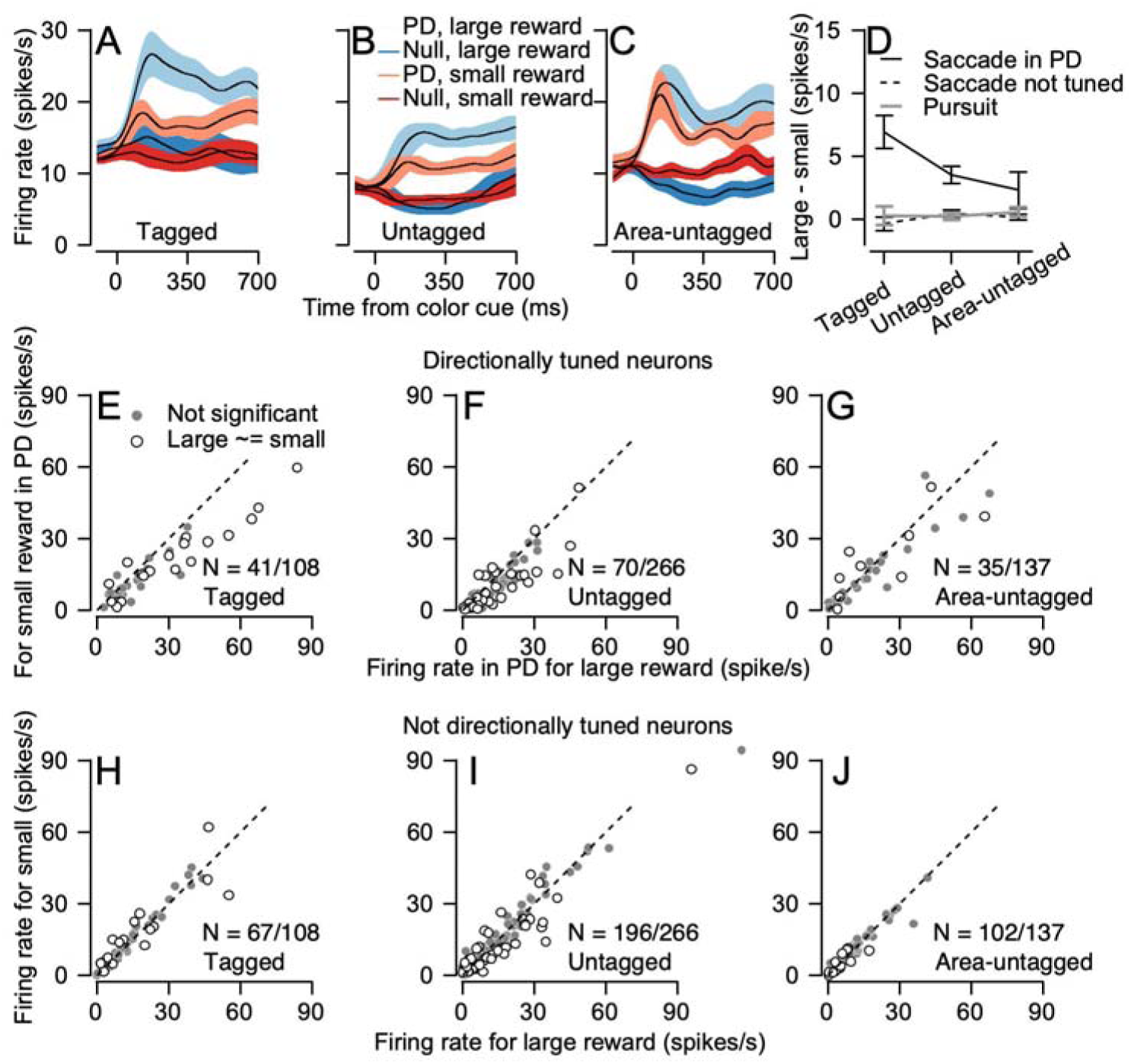
Reward modulation of directional tuning in the cue epoch. **A-C.** Average PSTH of the directionally tuned neurons for the three populations in the cue epoch of the saccade task. Light and dark colors show activity in the PD and the direction opposite to the PD (null). Blue and red show the activity in the large and small reward conditions. Bands show the SEM. **D.** Summary of the population average reward modulations (large - small). Solid line shows the modulation in the PD for neurons tuned to direction in the saccade task. Dashed line shows the average across all conditions for neurons that were not directionally tuned during saccade. Gray line shows the average activity for all neurons during pursuit (tuning was not tested experimentally). Error bars show the SEM. **E-J.** Scatter plot showing the reward modulations of single neurons during the cue. Values show the average activity between 0 and 500 ms after cue appearance for the large (horizontal) and small (vertical) reward conditions. Color of the spot indicates whether the difference between reward conditions was significant (open circles, p <0.05, Wilcoxon rank-sum test) or not significant (gray spots, p > 0.05). Different columns correspond to the three populations of neurons. The N in the plots is the number of the neurons shown in the plot out of the total number of neurons in each population. Top row (**E-G**) shows activity in the PD of neurons that were directionally tuned. Bottom row (**H-J**) shows the average activity across all directions for neurons that were not directionally tuned.

Reward did not consistently modulate the sign of activity in neurons that were not directionally tuned. Although reward had a significant effect in many of these neurons (70/365 open dots in Fig. 4H-J), the neurons that increased their activity in the large reward or small reward conditions were overall evenly distributed. As a result, the overall population difference between the large and small rewards did not reach significance (Fig. 4D, dashed line). Similarly, during the cue epoch of the pursuit task when directional information was not available, many neurons coded the reward (174/512) but the sign of the modulation was inconsistent across neurons (Fig. 4D gray line).

### Inconsistency of the sign of the reward modulation during movement

The analysis above indicated that in visually tuned neurons, reward potentiated directional tuning. We next tested whether this potentiation extended to movement. We first aligned the neural activity to the time of the saccade, to control for putative differences from the reward effect on saccade latency (Fig. 1C). In all populations, the overall directional tuning was only slightly modulated by reward (Fig. 5A-C) as indicated by the similarity of the population response in the PD at the time of the saccade. Neuron-by-neuron analysis confirmed that this lack of modulation was the result of the canceling out of opposite modulations of reward, as indicated by the distribution of neurons along both sides of the diagonal in Figure 5E-G. Similar averaging out was observed during pursuit (Fig. S5) and in neurons that were not directionally tuned during saccade and pursuit (Fig. 5D, H). These results extend our previous findings that focused on reward modulation during pursuit^15^ where we found a general inconsistency in the coding of reward and direction during movement in the FEF. This coding enables a readout of movement with common decoders such as the population vector, with minimal interference from reward^15^, thereby permitting simple population readouts that disentangle the multiplexed movement from reward information.

**Figure 5:**
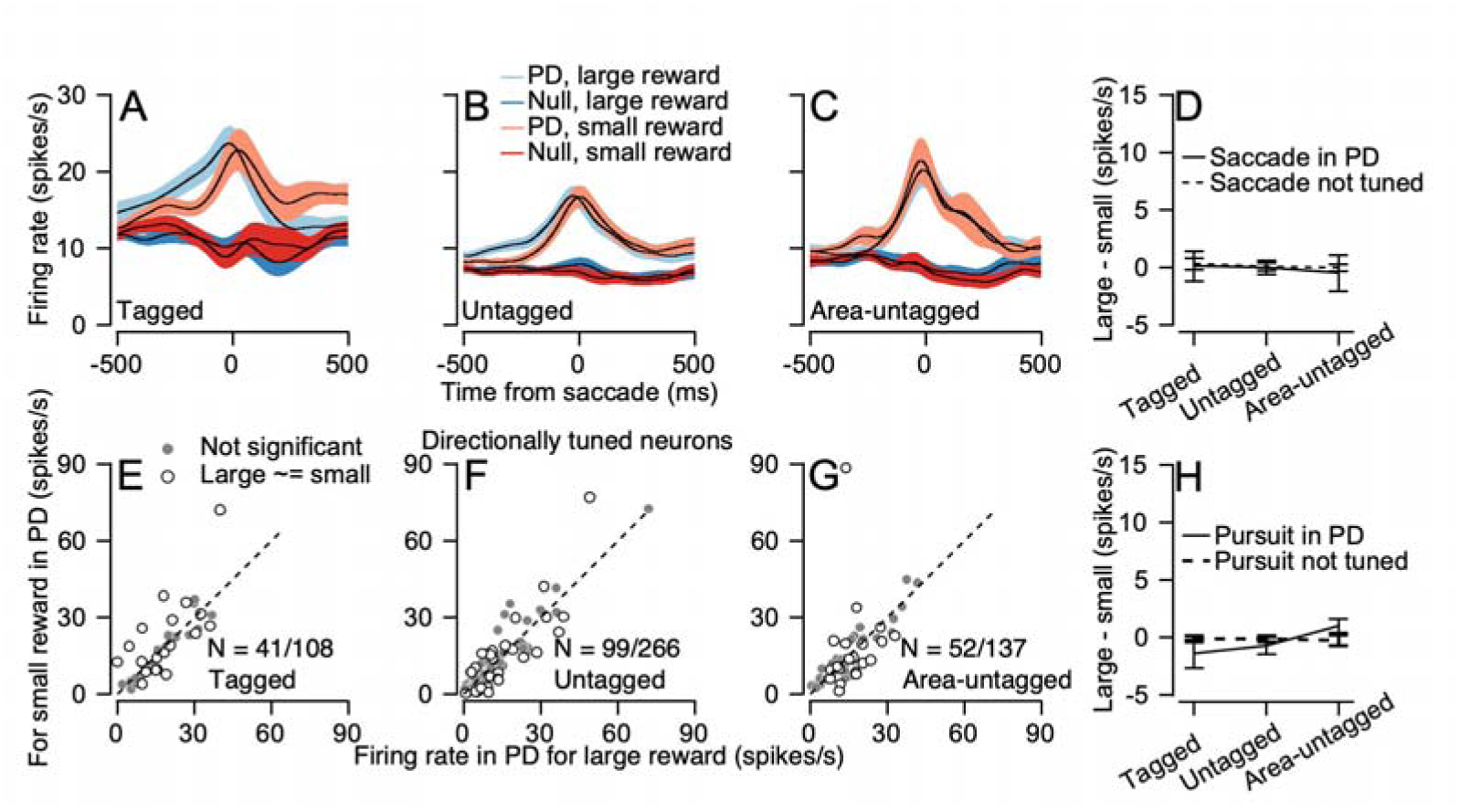
Reward modulation of directional tuning aligned to movement. **A-C.** Average PSTH of the directionally tuned neurons for the three populations aligned to the saccade. Light and dark colors show activity in the PD and the direction opposite to the PD (null). Blue and red show the activity in the large and small reward conditions. Bands show the SEM. **D, H.** Summary of the population average reward modulations (large - small) for the saccade (**D**) and pursuit task (**H**). Solid line shows the PD for neurons directionally tuned during the tasks. Dashed line shows the average across all conditions for neurons that were not directionally tuned during movement. Error bars show the SEM. **E-G.** Scatter plot showing the reward modulations of single neurons aligned to the saccade in the PD of directionally tuned neurons. Values show the average activity from 100 before to 300 ms after the saccade in the large (horizontal) and small (vertical) reward conditions. Color of the spot indicates whether the difference between reward conditions was significant (open circles, p <0.05, Wilcoxon rank-sum test) or not significant (gray spots, p > 0.05). Different columns correspond to the three populations of neurons. Only directionally tuned neurons are shown, to highlight the contrast of the results in this population with the cue results. The N in the plots is the number of the neurons shown in the plot out of the total number of neurons in each population.

### FEF inputs to the basal ganglia code the selected action before movement

Several theories have posited that the basal ganglia implement action selection^37^. The effect size analysis above indicated that in the choice task, the inputs to the basal ganglia had already differentiated between selection conditions. Specifically, activity was different if the large reward target appeared on the left or right side (dashed line in Fig. 2H). To further study neural representations during the choice task, we ran analyses comparing choice vs. single-target trials. We defined the *preferred choice condition* for each neuron as the choice condition with the largest response, and the condition with the smallest response as the *null choice condition*. In the choice task we contrasted opposite directions (left vs. right and up vs. down); thus, the *preferred choice condition* could differ from the PD in the tuning task which was based on more directions and conditions (see Methods). We then averaged the activity across neurons in the preferred and null choice conditions. We controlled for statistical biases by using half of the data from each neuron to determine the preferred choice condition and the other half to calculate the average activity and compared these to a single target. We found that soon after cue presentation, the FEF activity differentiated between choice conditions (Fig. 6A-C, solid versus dashed black line).

**Figure 6:**
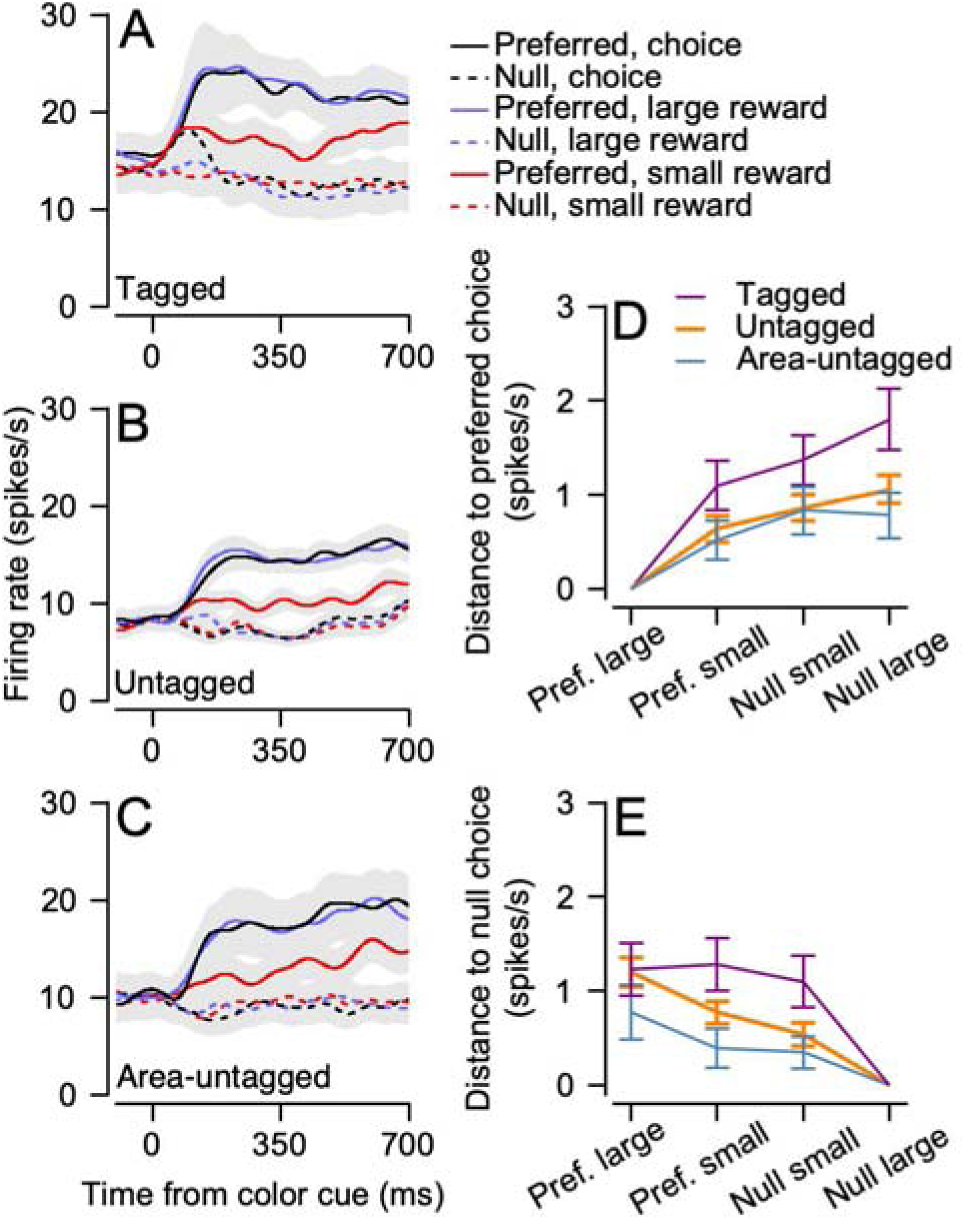
Choice signals in the FEF. **A-C.** Population average of neurons that significantly differentiated between choice conditions. Different colored lines correspond to the choice trials (black), large reward trials (blue) and small reward trial (red). Solid and dashed lines show activity in the preferred and null choice conditions. Different rows show different FEF populations. Gray bands show the SEM. **D, E.** The average distance (mean absolute value) between the choice in the preferred (D) or null (E) condition and single target conditions. To allow for paired comparisons the distance from choice to the corresponding single large condition was subtracted for each cell before the averages and SEM (bars) were calculated.

We refer to the single target conditions in which the monkey moved to the same location as the *preferred large and small reward conditions* and to the single target conditions in which the monkey moved in the null choice condition as the *null large and small reward conditions*. We then compared the activity during choice to these single target conditions (Fig. 6). In all populations the activity in the choice condition was almost identical to the activity in the corresponding large reward condition; i.e., activity in the choice trials, was similar to trials where the monkeys only observed the large reward target. This is shown by the similarity of the black and blue lines in the population average in Figure 6A-C and in the smallest PSTH distance between the preferred and null choice conditions corresponding to the large reward condition (Fig. 6D, E). This response in choice trials resembled a winner-take-all representation, in which the larger reward target is selected and the smaller reward is ignored.

This winner-take-all representation aligns with expectations associated with a selection process^38^ but not with other coding schemes. For instance, if activity in the choice conditions represented an average of the single-trial responses, we would expect the choice-preferred condition to be equidistant from the preferred large reward and the null small reward conditions. However, our results indicated that this was not the case (Fig. 6D). Another possibility is that activity during choice reflects reward potentiation of the visual receptive field. In this case, we would expect activity in the preferred choice condition to resemble the preferred large reward condition, since the larger reward target lies within the neuron’s receptive field. However, this interpretation also falls short. Specifically, in the null choice condition, where the small reward target is within the receptive field, we would expect activity to resemble the preferred small reward condition. Instead, the activity in the null choice condition more closely matched that of the null large reward condition (Fig. 6A-C, dashed black line resembles the dashed blue line rather than solid red and Fig. 6E). Thus, overall, the activity observed during choice was best interpreted as encoding the output of a selection process.

In the effect size analysis, we found slightly stronger coding of choice conditions in the tagged neurons, but this effect was not significant (dashed line in Fig. 2H). Therefore, in the analyses above, where we compared the choice and single target conditions, we did not include statistical comparisons between populations. However, we note that, as expected, the trends were consistent across analyses, with tagged neurons showing larger modulation.

## Discussion

We used optogenetics to identify corticostriatal neurons. We then examined the functional properties of these neurons in three eye movement tasks. Contrary to certain theories of the basal ganglia, we found that inputs to the basal ganglia already contained considerable information about the reward (Fig. 2, 3 and 4) and choice (Fig. 6). We also found an unexpectedly strong condition-independent modulation in the inputs to the basal ganglia (Fig. 2 and 3). Below we discuss the functional implications of these results.

### What is the role of the basal ganglia?

Reinforcement learning (RL) theories of the basal ganglia posit that the coding of reward prediction error by dopaminergic neurons^4,8,27,39–42^ serves as the instructive signal that potentiates cortical inputs associated with rewards. This potentiation could drive action selection^1–3^ or facilitate movement^43,44^. It is challenging to reconcile this model with our results, since a straightforward implementation would not predict that inputs from the frontal cortex already contain selection and reward information. Nevertheless, some adaptations of the RL model would remain consistent with our findings. For example, reward information in the FEF may originate from basal ganglia output projections to the cortex, a possibility which might be tested through lesions or latency comparisons^45^ of reward and choice signals in the cortex and the basal ganglia. Alternatively, the FEF-caudate network may not follow the RL model, while other basal ganglia subregions do, thus raising questions about the model’s generality. Finally, while initial learning may follow the RL model, later stages could involve a distributed representation across multiple areas, thus leaving the basal ganglia’s role beyond early learning uncertain.

Given that the RL model does not seem to explain our findings, what could the role of the basal ganglia be? Our current results and previous studies^30,31,46^ point towards a second type of model of the basal ganglia. These models suggest that rather than generating ‘content specific’ signals, the basal ganglia manipulate information to provide more concise or compact representations^47,48^. Algorithms such as dimensionality reduction could play an important role in reducing noise or reducing the physical space needed to store or read information. In a system that reduces dimensionally we would expect changes in the way neurons encode information rather than specificity in the coding of experimental variables.

Consistent with this supposition, in recent studies, we found that in the transformation from the input stages in the striatum to the basal ganglia output, the signal-to-noise of the responses to experimental variables increased^31^. This contrasts with a transformation that turns reward into movement signals, since the increase of the signal-to-noise was not limited to specific experimental variables. Furthermore, an analysis of the temporal profile of the basal ganglia output indicated that coding of the task variables by single neurons was high-dimensional^30^. This dimensionality was larger than the dimensionality of the striatal input as well as other populations in the eye movement system, including the sensory and frontal cortical areas, the cerebellar areas and the brainstem. Thus, our results, together with previous findings, suggest that the basal ganglia manipulate information rather than generate new representations. The causal role played by the basal ganglia in processing reward^49–52^ indicates that although reward might not be generated within the basal ganglia, the processing within the basal ganglia is critical.

### Interpreting the difference in effect size between groups

Ideally, we would want to split neurons into those that are connected to the basal ganglia and those that are not. However, untagged neurons cannot be classified as unconnected. The functional difference between the tagged and untagged neurons suggests that some of the unidentified neurons were unconnected. Therefore, the functional difference we observed between tagged and untagged neurons is likely to be an underestimation of the actual differences between neurons connected and unconnected to the basal ganglia.

The largest effect size for the tagged neurons indicates that the signal-to-noise of these neurons was the largest. This increase signal-to-noise could arise from the convergence of inputs from neurons with a similar signal, which would result in a sharpening of the response^31^. Thus, the difference in effect size between the tagged and nearby untagged neuron (Fig. 2, 3) suggests that in the transformation within the cortex, inputs with similar reward information or condition-independent modulations tend to converge on neurons that convey the cortical output to the basal ganglia.

The difference in effect size between the tagged and the area-untagged neurons could be due to several circuit motifs. One possibility is that the area-untagged neurons were located in FEF regions that are not directly connected to the basal ganglia and that these regions receive less reward and condition-independent information. A second possibility is that these neurons are part of the same microcircuitry as the other neurons but are in different layers. If this is the case, our results may point to an intra-layer computation^53,54^ in which reward information and condition-independent modulations are refined between cortical layers. Experimentally, it is difficult to distinguish between these possibilities, since the FEF lies within the arcuate sulcus, which makes it hard to identify specific layers using standard electrophysiological methods.

### Reward in the FEF

Our results confirm previous studies that have found reward and choice signals in the FEF^13–15,55,56^. On all tasks and in most task epochs, we found that many neurons differentiated between the small and large reward conditions. In the cue epoch of the saccade task, the reward-related activity potentiated directional tuning (Fig. 4). By contrast, in the movement epoch, the reward did not consistently modulate directional tuning (Fig. 4 and 5). This coding ensures that commonly suggested readouts of movement, such as population vector, can reliably read out the movement but are weakly modulated by reward. Similarly, population readouts that aggregate the single-neuron reward modulations^57^ average out movement-related activity. This organization would allow readouts of multiplexed signals during movement but not during the eccentric cue where reward consistently modulated the directional tuning (Fig. 4).

### Optogenetics in monkeys

Optogenetics is widely used in rodent research, but many studies on non-human primates continue to rely exclusively on electrophysiological methods. One major hurdle is that optogenetics in monkeys has been ineffective in driving movement. Our findings, along with other recent studies^26,58^, indicate that this limitation is not due to a lack of viral expression. We showed that optogenetics can be successfully applied to study neural pathways functionally, thus demonstrating its utility beyond causal manipulations. Our success in the expression of AAVretro in targeting cortico-striatal neurons in monkeys^59^ provides the technical foundation for studying the functional properties of the cortico-striatal neurons here. The high yield of neurons we identified suggests that this approach can be applied to studying other pathways as well. Thus, our findings make the case for expanding the use of optogenetics in non-human primates for functional circuit dissection.

## Methods

### Animals and ethics

We collected neural and behavioral data from two female (monkey K and monkey H) Macaca Fascicularis monkeys (4-5 kg). All procedures were approved in advance by the Institutional Animal Care and Use Committees of the Hebrew University of Jerusalem.

### Surgical procedures

We first implanted head holders to restrain the monkeys’ heads in the experiments. We performed a second surgery to place a round recording cylinder with an inner diameter of 19 mm over the FEF. The center of the cylinder was placed above the skull at 19 mm anterior and 15 mm lateral to the stereotaxic zero and tilted 25° with respect to the medial plane to allow for injections to the caudate nucleus. The localization of the FEF and caudate was based on stereotactic coordinates and then confirmed with post-implant MRI (Figs. S1 and S2) scan and electrophysiological markers (see below).

### Experimental setup

After the monkeys recovered from head holder surgery, they were trained to sit calmly in a primate chair (Crist Instruments) and consume liquid food rewards (baby food mixed with water and infant formula) from a tube set in front of them. We trained the monkeys to track spots of light that moved across a video monitor placed in front of them. Visual stimuli were displayed on a CRT monitor with a refresh rate of 85 Hz (55 cm from the eyes of the monkeys). The stimuli appeared on a dark background in a dimly lit room. A computer performed all real-time operations and controlled the sequences of target motions. The position of the eye was measured with a high temporal resolution camera (1 kHz, Eyelink 1000 plus, SR research) and collected for further analysis.

### Experimental design

The monkeys were engaged in saccade pursuit and choice tasks. In the saccade task each trial started with a bright white circular target that appeared in the center of the screen. After 500 ms of presentation, in which the monkey was required to acquire fixation, a colored target appeared at a position 10° eccentric to the fixation target in one of the four cardinal directions. We defined this time point as the cue onset. The color of the target indicated that upon successful completion of the trial the monkey would receive either a small or large reward (0.07 and 0.15 ml of food). For monkey K we used blue and red targets for the large and small rewards and for monkey H we reversed the associations. The monkeys were required to maintain fixation on the central target, and moving the eye towards the eccentric target would abort the trial. After a variable delay of 500-700 ms, the fixation target disappeared which served as the Go signal for the monkey to move their eyes to the peripheral target. The monkeys had to acquire fixation in 750 ms and then after an additional 200-300 ms they received the reward.

The structure of the pursuit trials was similar to the saccade trials but with the following differences. A colored target replaced the white target in the center of the screen. After the cue epoch the target jumped 4° in one of the four cardinal direction and then moved in the opposite direction towards and through the center of the screen at 20°/s (step-ramp^60^). The target moved for 750 ms, then stopped and stayed still for an additional 200-300 ms.

The structure of the choice trials was also similar to the saccade task but instead of a single target, two colored eccentric targets appeared at the cue. The target appeared either along the horizontal or vertical axis 10° eccentric to the fixation target. When the central target disappeared, the monkey was free to select the target to which to saccade. Online we detected the saccade as an eye movement that exceeded 80°/s. The target that was closer to the eye at the end of the saccade remained and the other target disappeared. The monkeys were then required to continue to fixate on this target and received a reward based on the color of the target. In all tasks we enforced a precision window of 3-5°x3-5° around the fixation target with grace periods after the Go signal to allow the monkeys to acquire the target. Trials in which the monkeys failed to meet the fixation requirements were removed from the analysis. In each recording session we interleaved saccade, pursuit and choice trials such that they contained 80 pursuit and 80 saccade trials (4 directions X 2 reward conditions X10 repetitions), and 80 selection trials (2 directions X 2 target configuration conditions X 20 repetitions).

### Virus injection

We injected AAVretro-hSyn-ChR2-GFP virus (ELSC virus core) to the striatum. We first injected the virus to the striatum of mice and confirmed the expression with histology and recorded LFP and single units in the cortex in responses to light stimulation. We then injected the same virus to the monkeys. We made efforts to confirm the injection site without histology, as we aim to release the monkeys to a sanctuary at the end of the experiments. Before the injections we mapped the recording chamber, and then lowered single electrodes (Thomas recordings Mini matrix) to identify the striatum. Identification was based on well-known physiological and anatomical characteristics including the depth of the electrode compared to the MRI scan and atlas, the shape of the extracellular spike associated with medium spiny neurons (MSN), the characteristic low firing rate of the MSN, the presence tonically active neurons (TANs) with their broad extracellular spike and a gap in the recorded neurons after penetration of the cerebral cortex.

After a few weeks of mapping, we injected the virus. We loaded low-viscosity silicone fluid (polydimethylsiloxane, viscosity: 1 cSt at 25°C, Merck) into a 1 ml syringe and a thick-walled plastic tubing (inner diameter: 0.25 mm, length: 50 cm, Thomas Recordings injection system). The fluid fluoresced in color to allow us to identify the boundary between the silicon and the virus to confirm the injections. After loading the virus, we lowered a recording electrode and an aligned injection micropipette (Thomas Recordings, 35 gauge, lowered with Mini matrix). Based on the extracellular activity we identified the upper edge of the striatum. We then searched for the lower edge of the striatum and then injected 1-2 mm above the lower edge to 1-2 mm before the upper edge of the striatum 1 mm apart. In each site we injected 2-3 μl of the virus (titer 2.3*10^12^) at a speed of 15nl/s. We then waited 5 min before we pulled the electrode 1 mm upwards to the next site. In the upper site of the striatum, we waited 30 min before extracting the electrode from the brain. In each penetration we injected 1-4 sites depending on the thickness of the striatum. We aimed the injections towards the eye movement areas in the caudate (Fig. S1). Overall, we injected in 10 and 9 penetrations in monkeys K and H.

To further verify the injection location, for several injections we added Manganese (Mn^+2^, 0.01M) to the injected solution. Then, a few hours after the injection, we scanned the monkeys in an MRI using contrasts that make Mn^+2^ visible (Fig. S1, sequence details: T1, flash, TR=500, TE=3.7, FA=30). In the first monkey we performed the scan in which we injected only Mn^+2^ to validate our injection system before injecting the virus. After validation in the second monkey, we added Mn^+2^ to the same solution we used to inject the virus.

### Neural recordings and spike sorting

On each recording day, we lowered an optrode (an optical fiber connected to a tetrode, OD=356 μm, Thomas Recordings) to the FEF. When we reached the site with the neurons, we applied light stimulation (LED light, wavelength: 470nm) protocol that included 10 repetitions of 10 pulses (5 ms durations) of stimulation at 10, 20 and 50 Hz (overall 10X10X3 = 300 pulses). Online we monitored the activity and searched for responses of single units or local field potential (LFP) to the light stimulation. When we observed a response, we applied the behavioral protocol. Often, we found a relatively thin layer (typically less than 2 mm, Fig. S2) in which neurons or the LFP responded to the stimulation, in contrast to widespread activity that would be expected if the virus had leaked to the recording site. For a more complete comparison we also recorded neural activity in areas where we did not observe responses to the stimulation. We sorted the spikes offline (Plexon offline sorter) and only well-isolated units were included in the analysis.

To confirm that the neurons that responded to light stimulation were the same neurons that were recorded during the task, we calculated the correlation and distance between the spike waveform of all the cells that recorded during the task session and the light-evoked response waveform (Fig. S3). 86% (93/108) of the neurons identified as connected met a previously used criteria for ensuring that light-evoked waveforms were similar to the task waveform (correlation coefficient > 0.9)^27^.

### Quantitative and statistical analyses

To detect responses to the optical stimulation we calculated the spike rate in the 20 ms after the stimulation in 1 ms bins (in units of spikes/ms). We then subtracted the rate before the stimulation from the rate after the stimulation to calculate the rate of increase of the spikes in each ms after the stimulation. We excluded the 50 Hz stimulation condition in this analysis since the inter-stimulus-interval was too short to establish the pre-stimulus activity. Neurons were characterized as responding to the stimulation if the cumulative distribution of the increase in rate exceeded 0.5. This criterion corresponds to an additional spike above baseline within 20 ms of stimulus onset in 50% of the trials. and is much more stringent than a statistical test for an increase in rate. We used this stringent criterion since even a small number of errors in the spike sorting in which a non-responding neuron was contaminated with a responding neuron could lead to a significant increase. Finally, we confirmed that only neurons were included in which the response had a latency of less than 5 ms as determined by an increase in rate of at least 0.2 in the first 5 ms after the stimulation. This additional screening only removed two neurons and was practically negligible. The criterion was conservative in comparison to the classification of a human observer. All the neurons we classified as responding were also classified as responding by a human observer (N=108), but the human observer classified 57 other neurons as responding. Repeating the analysis based on a human observer classification did not alter our conclusions. Overall, 73, 163 and 50 neurons were classified as tagged, untagged and area-untagged in monkey H and 35, 103 and 88 were classified as tagged, untagged and area-untagged in monkey K.

To calculate the coding of a variable throughout an epoch, we calculated the partial *ω*^2^ in ANOVA models that included time, direction and reward as variables. *ω*^2^ is a common effect size measure used in ANOVA designs that is often preferred over other effect size measures since it is unbiased. We calculated the number of spikes in 250 ms bins, and then calculated the *ω*^2^ as follows:

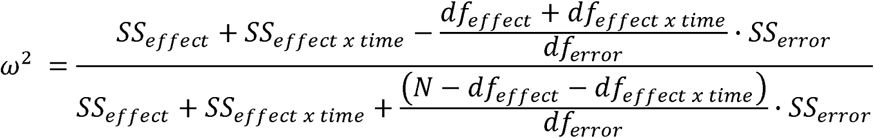

where *SS_effect_* is the ANOVA sum of squares for the effect of a specific variable, *SS_effect x time_* is the ANOVA type II sum of squares for the interaction of a specific variable with time, *SS_error_* is the sum of squares of the errors after accounting for all experimental variables, *df_effect_*, *df_effect x time_* and *SS_error_* are the degrees of freedom for a variable, the interaction of the variable with time and the error, respectively. We included the interaction term since it quantifies the time-varying coding of the variable. *N* is equal to the number of trials x the number of time bins. To calculate the time effect size, we did not include interaction term or the corresponding degree of freedom.

For the cue and outcome epochs we used a time window from 250 before to 500 ms after the event onset. For movement, we used a time window of 0-1000 ms after the Go signal. Changing the bin size or the time window yielded highly correlated effect sizes. The direction variable was not in the ANOVA design in the pursuit cue epoch. The reward size was not in the ANOVA design in the choice task. In all other epochs, reward, direction and time were included as variables. We also quantified all possible interactions (Reward x Direction, Reward x Direction x Time) but these did not yield any consistent results and were often small in comparison to other values. We used the partial effect size since it enables a better comparison between neurons that responded or did not respond to other experimental variables. Using the full effect size did not alter any of our conclusions. To test for differences between the average effect size across populations we used a permutation test. We shuffled the group labels, and calculated the difference between the average effect size of the shuffled populations. We repeated this 1000 times and if the actual average difference was larger than 95% of the shuffles the difference was considered significant.

To determine the preferred direction (PD) of each neuron, we calculated the direction of the center of mass of the responses and assigned the PD to the cardinal direction that was closest to the calculated direction. We used both large and small reward trials to calculate a single PD for each cell in each task and task epoch. We used a window of 50-500 ms after the cue. When aligning the saccade, we used a window from 100 ms before to 300 ms after the saccade. We used a shorter time before the saccade to minimize the effect of the cue on the tuning of the saccade. For pursuit we used a window of 50-500 ms after motion onset. We confirmed that the results were insensitive to the time windows. We used these same time windows to determine whether a neuron responded significantly to the reward or directions. In each trial we counted the number of spikes in the window and performed a 1-way (Direction or reward) or a 2-way (Direction and reward) ANOVA to test for significance of the response to a variable.

We calculated the peristimulus time histogram in bins of 1ms and smoothed with a 30 ms SD Gaussian. We calculated the population PSTH by averaging the responses from all neurons in the PD or Null direction. Since we were interested in the effect of the reward on tuning and we used both rewards for calculating the PD therefore the effect of reward on tuning was not biased by this analysis. To avoid bias in the analysis of choice conditions, we used half of the choice trials to test whether a neuron’s response was significantly different across the choice conditions and defined the preferred choice direction based on which direction had the larger firing rate. We then used the other half of the trials to plot the population PSTHs and perform comparisons with the single target trials. This procedure insured that the population responses were not statistically biased by the selection of the preferred direction.

### Declaration of generative AI and AI-assisted technologies in the writing process

During the preparation of this work the authors used ChatGPT in order to edit English. After using this tool/service, the authors reviewed and edited the content as needed and take full responsibility for the content of the publication.

## Acknowledgments

This project received funding from the European Research Council (ERC) under the European Union’s Horizon 2020 research and innovation program (grant agreement No. 755745).

