## Supplemental Figures 1-5 for "Enhanced reward coding and condition-independent dynamics in optogenetically identified corticostriatal neurons in monkeys"

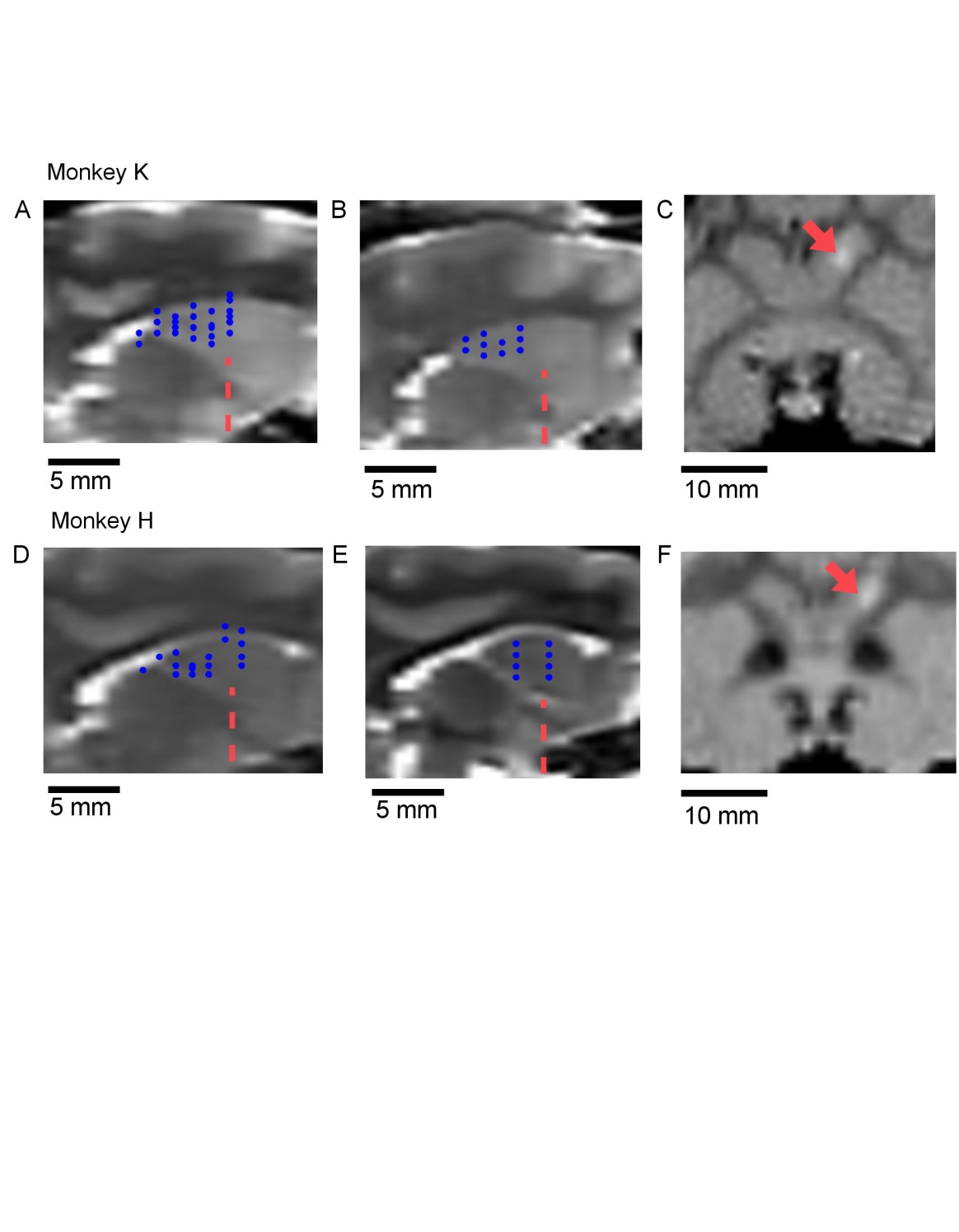

**Figure S1: Virus injected sites aligned to the MRI and mn+2 verification. A,B,D,E:** Sagittal sections 3 (**A,D**), 2 (**B)** and 4 **(E**) mm lateral to the mid-sagittal section. Blue dots mark the estimated site of injection of the virus. Dashed red line shows the anterior commissure coronal plane (AC 0). **C, F:** Examples of coronal sections of an MRI from days in which we injected Mn+2 before the MRI. Sections are at (**C**) and 2 mm posterior (**F**) to the anterior commissure. The arrow point towards the Mn+2. Top and bottom rows show MRI for monkey K and H. Calibration bars show 5 (**A,B,D,E**) and 10 (**C,F**) mm**.**

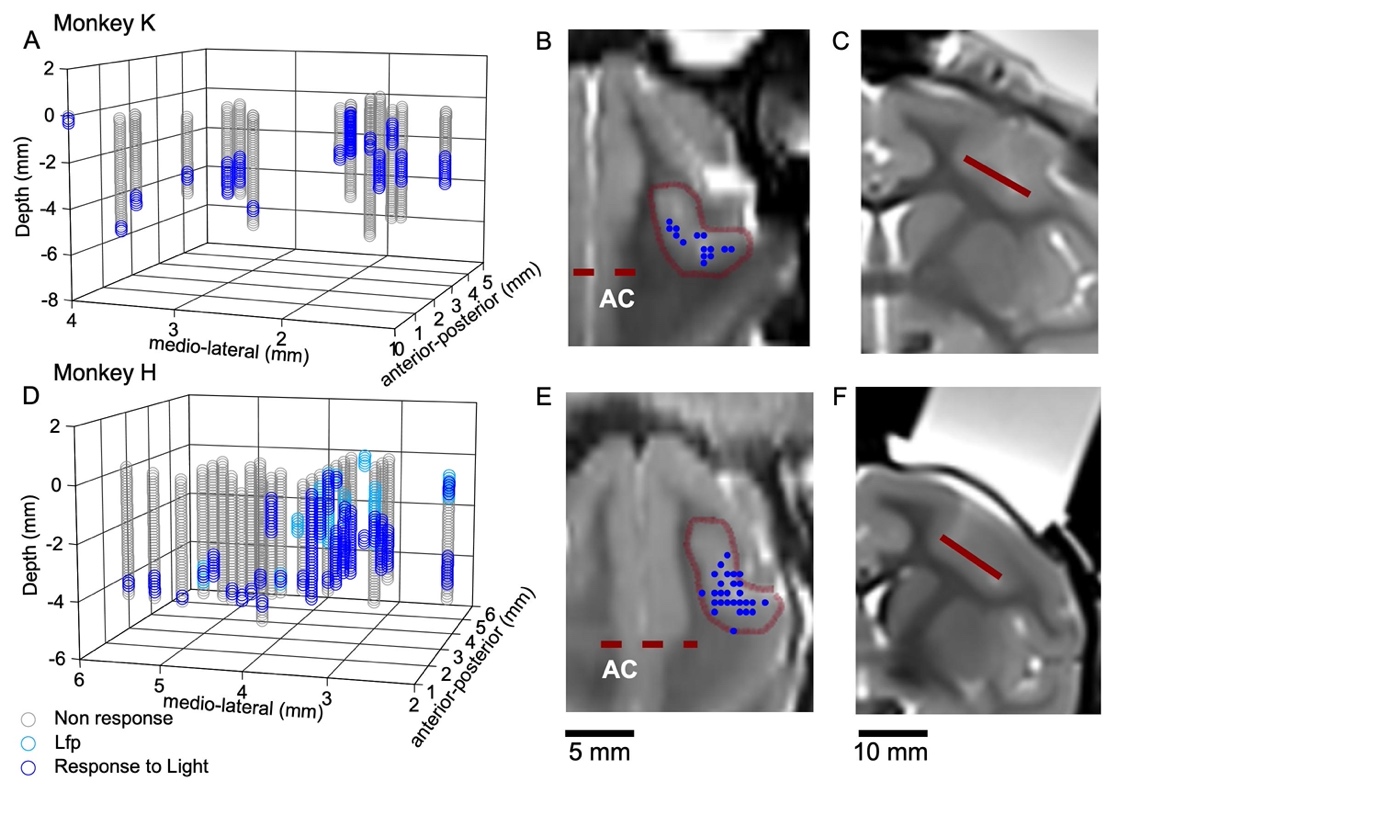

**Figure S2: Estimated recording sites in the FEF. A, D.** Three-dimensional plot showing the location of the recorded neurons in coordinates of the recording chamber. The chamber was placed above the FEF along the anterior-posterior axis and tilted 25° from the sagittal plane. The axes show the distance along the medio-lateral, anterior-posterior and depth (z) in the chamber. Gray markers show locations in which we did not find responses to the light stimulation. Dark blue markers show sites in which neurons responded to the light stimulation. Light blue markers show sites in which we found a response to light in the local field potentials (LFP) but not in the spikes of the neurons. **B,C,E,F:** Registration of the three dimensional chamber map onto the MRI**. B, E:** A section through the MRI showing the location of the recording in blue dots. Red line shows an estimate of the arcuate sulcus based on the MRI contrast. The section is parallel to the chamber surface and perpendicular to the motion of the electrode. Dashed red line shows the anterior commissure plane. **C, F:** Coronal section at the level of the anterior commissure showing in red the depth of the section shown in **B** and **E**. Calibration bars show 5 (**B,E**) and 10 (**C,F**) mm.

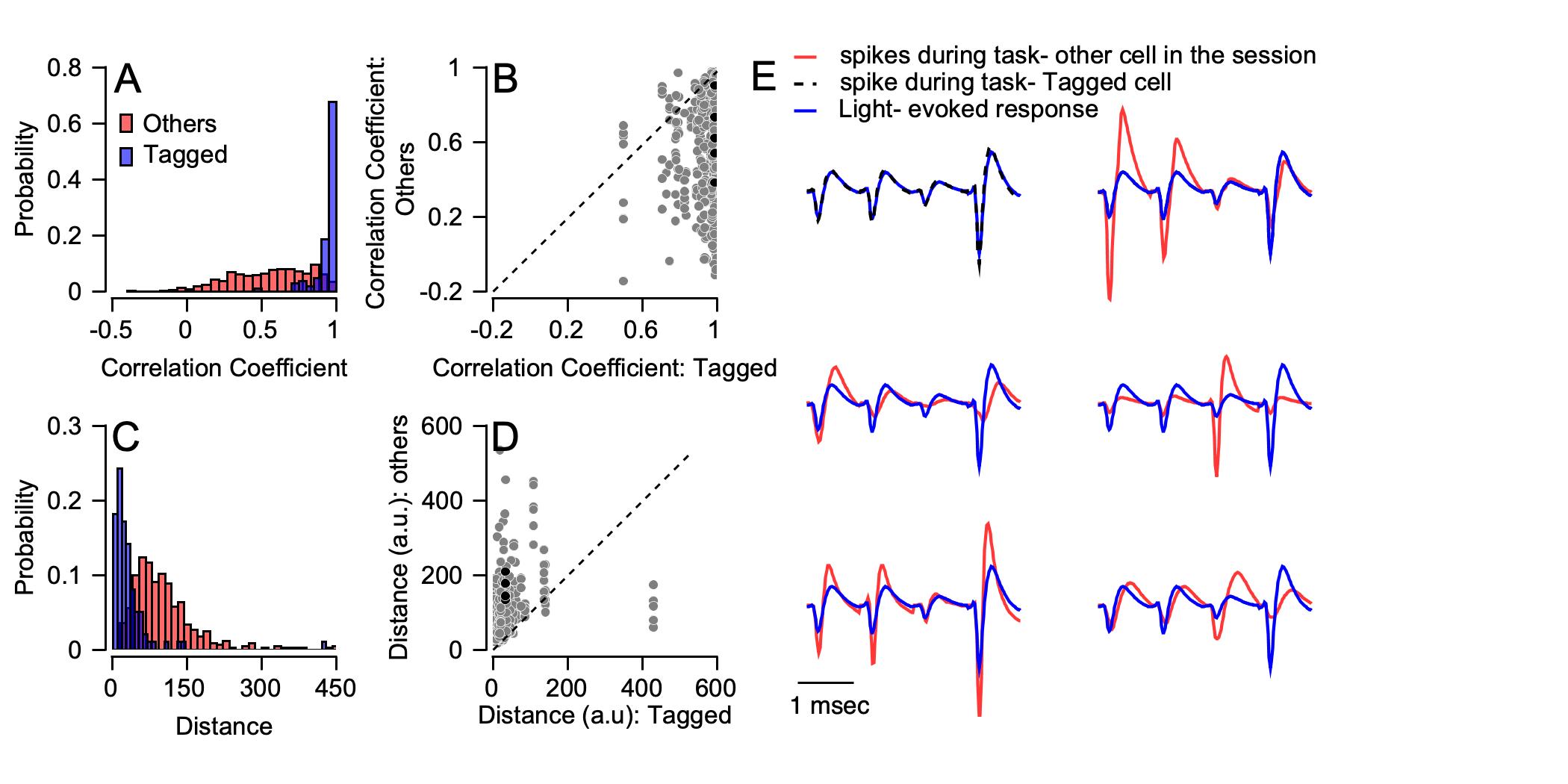

**Figure S3: Analysis of extracellular waveforms. A, D.** Distribution of the correlation coefficient (**A**) and Euclidean distances (**D**) between the average tetrode waveforms during optical stimulation and the behavioral task. The blue histogram shows the distribution of values calculated for connected neurons with themselves and the red histogram shows the distribution of values calculated for connected neurons during stimulation with other simultaneously recorded neurons during the task. **B, D**. Neuron-by-neuron comparison of the correlation coefficient (**B**) and distance (**D**) for the connected neurons with themselves (Horizontal) and the connected neurons with other neurons recorded simultaneously (vertical). **E**. Example of the waveform of the light- evoked response of the connected neuron (blue) and the response of the same neuron during the task (black) and other neurons during the task (red). Black dots in **B** and **D** show the correlation coefficient and distance for the example neuron.

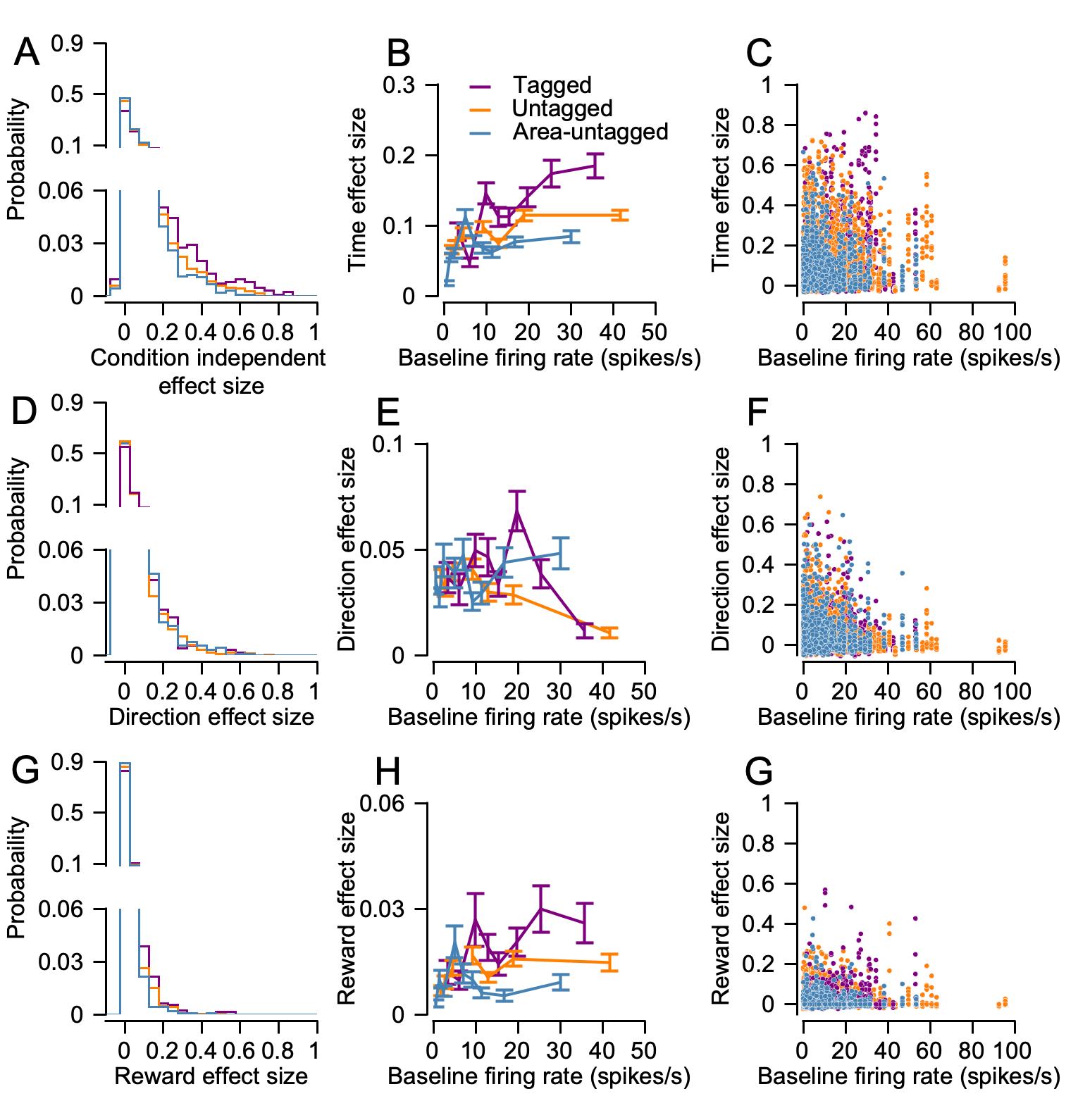

**Figure S4: Effect size analysis controlled for baseline firing rate. A,D,G.** Full distribution of the time (**A**), direction (**D**) and reward (**G**) effect sizes, data were gathered across all relevant tasks and task epochs. Histogram were split in the middle and the vertical axis adjusted to emphasize the tail of the distribution. **B,C,E,F,H,I.** Effect size as a function of the baseline firing rate. In the left column (**C, F, I**), each dot represents the effect size for one task and task epoch, plotted as a function of the overall pre-cue average. The middle column (**C,E,H**) shows the average and the SEM of the effect size in 10 bins with equal numbers of data points split by the baseline rate.

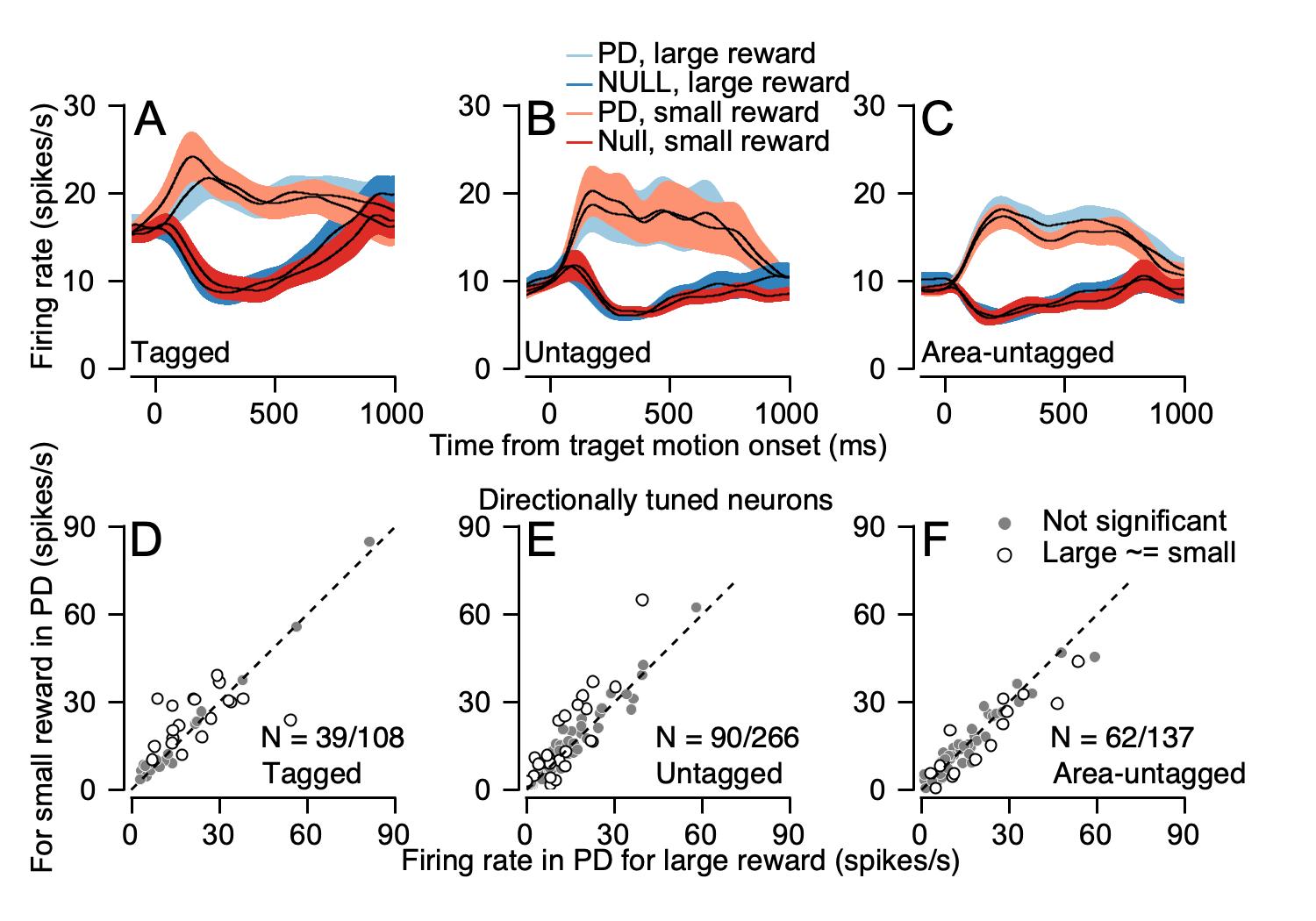

**Figure S5: Reward modulation of directional tuning during smooth pursuit movement. A-C.** Average PSTH of the directionally tuned neurons for the three populations aligned to motion onset. Light and dark colors show activity in the PD and the direction opposite to the PD (null). Blue and red show the activity in the large and small reward conditions. Bands show SEM. **D-F.** Scatter plot showing the reward modulations of single neurons in the PD of directionally tuned neurons. Values show the average activity during the target motion in the large (horizontal) and small (vertical) reward conditions. The color of each spot indicates whether the difference between reward conditions was significant (open circles, p < 0.05, Wilcoxon rank-sum test) or not significant (gray dots, p > 0.05). Different columns correspond to the three populations of neurons. The N in the plots is the number of the neurons shown in the plot out of the total number of neurons in each population.
